# Brain structure-function coupling provides signatures for task decoding and individual fingerprinting

**DOI:** 10.1101/2021.04.19.440314

**Authors:** Alessandra Griffa, Enrico Amico, Raphaël Liégeois, Dimitri Van De Ville, Maria Giulia Preti

## Abstract

*Brain signatures* of functional activity have shown promising results in both *decoding* brain states, meaning distinguishing between different tasks, and *fingerprinting*, that is identifying individuals within a large group. Importantly, these brain signatures do not account for the underlying brain anatomy on which brain function takes place. Structure-function coupling based on graph signal processing (GSP) has recently revealed a meaningful spatial gradient from unimodal to transmodal regions, on average in healthy subjects during resting-state. Here, we explore the potential of GSP to introduce new imaging-based biomarkers to characterize tasks and individuals. We used multimodal magnetic resonance imaging of 100 unrelated healthy subjects from the Human Connectome Project both during rest and seven different tasks and adopted a support vector machine classification approach for both *decoding* and *fingerprinting*, with various cross-validation settings. We found that structurefunction coupling measures allow accurate classifications for both task decoding and fingerprinting. In particular, key information for fingerprinting is found in the more liberal portion of functional signals, that is the one decoupled from structure. A network mainly involving cortico-subcortical connections showed the strongest correlation with cognitive traits, assessed with partial least square analysis, corroborating its relevance for fingerprinting. By introducing a new perspective on GSP-based signal filtering and FC decomposition, these results show that brain structure-function coupling provides a new class of signatures of cognition and individual brain organization at rest and during tasks. Further, they provide insights on clarifying the role of low and high spatial frequencies of the structural connectome, leading to new understanding of where key structure-function information for characterizing individuals can be found across the structural connectome graph spectrum.

**Highlights:**

- The relation of brain function with the underlying structural wiring is complex
- We propose new structure-informed graph signal processing (GSP) of functional data
- GSP-derived features allow accurate task decoding and individual fingerprinting
- Functional connectivity from filtered data is more unique to subject and cognition
- The role of structurally aligned and liberal graph frequencies is elucidated

## 1. Introduction

The existence of *brain signatures* based on functional magnetic resonance imaging (fMRI), meaning specific features uniquely characterizing either tasks or individuals, has emerged from the development of advanced data analysis methods in the last two decades. On the one hand, the application of pattern recognition techniques to neuroimaging data proved the capability of fMRI to *decode* task-specific brain activity (Gao et al., 2020; Haynes and Rees, 2006; Li and Fan, 2019; Richiardi et al., 2011; Wang et al., 2020). Significant progress in this direction was made by the recent advent of deep learning (Gao et al., 2020; Li and Fan, 2019; Wang et al., 2020), even if it remains nontrivial to interpret the biological meaning of the learned features. On the other hand, similarly to a *fingerprint*, fMRI-based features can accurately identify individuals from a large group (Amico and Goñi, 2018a; Biazoli et al., 2017; Finn et al., 2015; Mansour L et al., 2021; Van De Ville et al., 2021). In a seminal paper by Finn and colleagues (Finn et al., 2015), functional connectivity (FC) profiles were used to successfully classify subjects across resting state test-retest sessions, and even between task and rest conditions. The fronto-parietal network emerged as the main contributor to subject discrimination, and was shown to predict individual cognitive behavior (i.e., level of fluid intelligence). In addition to functional activity, brain anatomical features, such as cortical morphology and white-matter structural connectivity, were also proven useful for brain fingerprinting (Kumar et al., 2017; Lin et al., 2020; Valizadeh et al., 2018; Wachinger et al., 2015; Yeh et al., 2016).

In this context, a still unexplored brain feature, which could offer new insights into task decoding, individual fingerprinting and behavioural correlates, is the degree of coupling between function and structure, i.e., how brain functional activity and connectivity align to the underlying structural connectivity architecture as measured with diffusion-weighted (DW) MRI. Early attempts to investigate structure-function relationships in the brain spanned from simple approaches, such as correlational analyses (Amico and Goñi, 2018b; Goñi et al., 2014; Honey et al., 2009; Mišić et al., 2016; Zhang et al., 2011), to more complex ones, like whole brain computational and communication models (Amico et al., 2021; Avena-Koenigsberger et al., 2018; Deco et al., 2011; Griffa et al., 2017; Mišić et al., 2015; Seguin et al., 2020). More recently, graph signal processing provided a novel framework for a combined structurefunction analysis (Huang et al., 2018; Medaglia et al., 2018; Preti and Van De Ville, 2019). Within this setting, Preti and Van De Ville quantified the degree of structure-function dependency for each brain region, by means of the newly introduced Structural-Decoupling Index (SDI) (Preti and Van De Ville, 2019). This nodal metric quantifies the degree of local (dis)alignment between structure and function, and it is obtained by decomposing the structural connectome into harmonics in the graph frequency domain, and projecting the functional signals (fMRI frames at each timepoint) in the space spanned by the structural harmonics. The functional signals are then filtered into low and high structural graph frequencies, giving rise to coupled and decoupled signal components, respectively. The ratio between the energy of these two signal portions yields the SDI of a brain region. During resting state in healthy subjects, local structure-function (de)coupling showed a very characteristic and behaviorally relevant spatial distribution, spanning from lower-order functional areas such as visual and somatosensory cortices, with activity highly constrained by the structure underneath, to higher-order ones, with activity more liberal. However, the extent to which this configuration changes in different task-related states, or in different subjects, still remains unexplored. Moreover, the quantification of the structure-function coupling at the level of single brain connections may bring new insights into brain organization principles and their uniqueness to brain states and individuals. In particular, do structure-function dependency patterns represent a signature of a particular task-related state? Can they act as a brain fingerprint uniquely identifying individuals? And which structure-function dependency features are more relevant to task decoding, subject fingerprinting, and inter-individual cognitive variability?

To answer these open questions, we analysed the structural and functional data during resting state and seven different tasks of 100 unrelated healthy subjects from the Human Connectome Project (HCP) (Van Essen et al., 2013), and obtained their structure-function signatures quantified through: (i) the SDI, and (ii) a new GSP-based decomposition of the FC. The latter is obtained by assessing the functional connectivity between fMRI signal components that are more coupled or decoupled to the underlying structure, named coupled-FC (c-FC) and decoupled-FC (d-FC), respectively. These GSP-derived features quantify brain structurefunction coupling at the level of either single brain regions (SDI) or single brain connections (c-FC, d-FC) and were used to classify different tasks and individuals. In both cases, the classification showed high accuracy for all the three structure-function coupling measures, across various cross-validation settings. Two specific networks including regions that are key to either *task decoding* or *individual fingerprinting* based on structure-function coupling emerged. Results were then compared with the classification performances obtained with conventional nodal (node strength) and edgewise measures of FC, without knowledge from the underlying structure. Finally, nodewise and edgewise structure-function couplings in resting state were shown to correlate with individual cognitive traits including fluid intelligence and sustained attention, particularly in the high-frequency FC components (d-FC) of the structural connectome.

## 2. Material and Methods

### 2.1 Methods Outline

The methodological pipeline is illustrated in Fig. 1. From the fMRI time courses (Fig. 1A) of 100 individuals during rest and seven tasks, conventional edgewise and nodal FC measures (FC matrix and FC node strength) were computed (Fig. 1B). In parallel to that, the GSP pipeline outlined in (Preti and Van De Ville, 2019) was implemented to decompose functional signals at each timepoint onto the underlying structural bases and filter them in coupled (low-frequency) and decoupled (high-frequency) portions (Fig. 1C). Structure-function coupling was then evaluated at the level of connections and regions, by means of coupled and decoupled FC and structural-decoupling indexes, respectively (Fig. 1D). c-FC and d-FC are FC matrices derived from the coupled and decoupled portions of fMRI time courses. The SDI quantifies instead the amount of local alignment between brain functional signals and the underlying structural connectivity network at the nodal level. Next, the task decoding and individual fingerprinting accuracy obtained from the nodal and edgewise structure-function coupling features (SDI, c-FC and d-FC, Fig. 1D), as well as from corresponding nodal and edgewise measures of FC not taking into account the underlying brain structure (Fig. 1B), were assessed with Support Vector Machine (SVM) classification (Fig. 1E) and compared. Finally, multivariate relationships between the different nodal and edgewise features and individual cognitive traits were assessed with partial least square correlation (PLSC) analyses.

**Figure 1.**
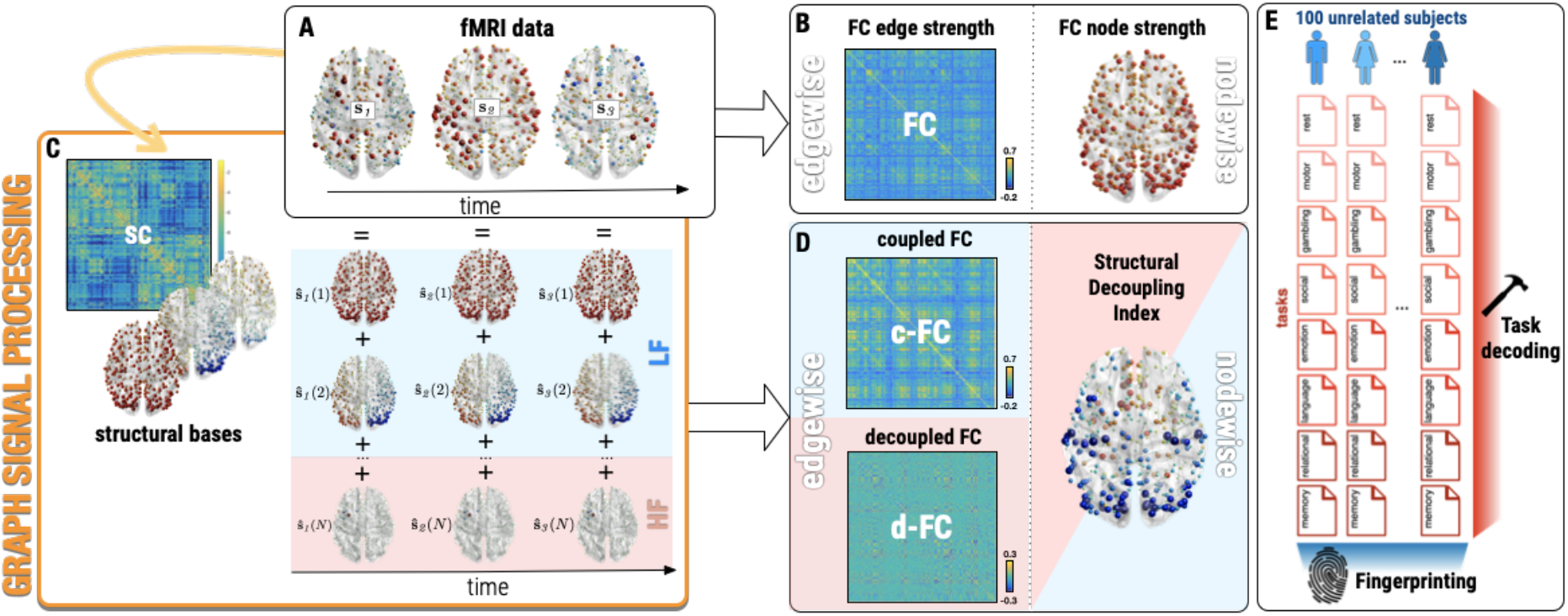
Method workflow. From fMRI nodal signals at each timepoint (A), functional connectivity (FC) is evaluated through conventional edgewise (FC matrix) and nodewise (FC node strength) measures (B). The graph signal processing (GSP) pipeline is applied to decompose functional signals into the structural harmonics obtained from the eigendecomposition of the structural connectome (SC) Laplacian (C). Functional signals are then filtered into two components; i.e., one coupled and one decoupled from structure, by applying ideal low pass (light blue) and high pass (pink) filters in the graph spectral domain (C). Edgewise and nodewise metrics evaluating structure-function coupling are obtained by computing FC matrices from coupled and decoupled signals (coupled and decoupled FC (c-FC and d-FC), respectively), and the structural decoupling index (SDI). Edgewise and nodal measures of both FC (B) and structure-function coupling (D) enter separate support vector machine (SVM) classifications with various cross validation settings to test their task decoding and fingerprinting value, quantified by task and subject identification accuracies (E).

### 2.2 Data & Preprocessing

The *N_S_* = 100 unrelated healthy subjects of the HCP dataset U100 - HCP900 data release (54 females, 64 males, mean age = 29.1 ± 3.7 years) were included in the study. Ethical approval was obtained within the HCP. Analyses were restricted to these 100 subjects to ensure absence of any family relationship which may influence fingerprinting results. fMRI acquired with *N_T_* = 8 different task conditions (resting state and 7 tasks: emotion, gambling, language, motor, relation, social, working memory), each recorded with *N_E_* = 2 phase encoding directions (right-left and left-right), as well as DW-MRI sequences were pre-processed with state-of-the-art pipelines, in order to obtain regional functional time courses and their structural connections, based on a parcellation with *N_ROI_* = 379 regions (360 cortical areas (Glasser et al., 2016) and 19 subcortical ones as provided by the HCP release (Fischl et al., 2002; Glasser et al., 2013)). Each cortical area was assigned to one of the 7 Yeo networks through majority voting procedure for *post hoc* analyses (Yeo et al., 2011). Minimally preprocessed data from the HCP were selected (Glasser et al., 2013; Van Essen et al., 2013) and the following additional pre-processing steps were performed. Nuisance signals were removed from voxel fMRI time courses (linear and quadratic trends, six motion parameters and their first derivatives, average white matter and cerebrospinal fluid signals and their first derivatives) and average time courses were computed in each region of the parcellation, previously resampled to the functional resolution, and z-scored. To remove the effect of the paradigm on task data, only for task classification, paradigms were regressed out trial by trial from functional time courses (a separate regressor for each task trial was included in the model). Functional connectomes were obtained as Pearson’s correlation between pairwise time courses and FC nodal strength was computed for each region as the sum of absolute values of all the connections of that region (Fig. 1B).

The same DW-MRI processing pipeline detailed in (Preti and Van De Ville, 2019) was used to reconstruct whole brain tractograms including 2 million fibers, using a spherical deconvolution approach and the Spherical-deconvolution Informed Filtering of Tractograms 2 (SIFT2 (Smith et al., 2015), https://www.mrtrix.org/). Structural connectomes were then obtained, after resampling of the same parcellation to diffusion space, as the number of tracts connecting two regions, normalized by the sum of the two regions’ volumes. An average structural connectivity (SC) matrix, representative of the whole population, was obtained by averaging the structural connectivity values across subjects.

### 2.3 Structure-function coupling features

The graph signal processing framework detailed in (Preti and Van De Ville, 2019) was adopted to obtain structure-function signatures (the SDI and the newly introduced c-FC and d-FC) for each subject and acquisition. In brief, the average SC across the population is decomposed into structural harmonics *u_k_* by eigendecomposition of the SC Laplacian *L* = *I* – *A_symm_* (given the identity matrix *I* and the symmetrically normalized adjacency matrix *A_symm_* of the SC):

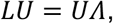

where each eigenvalue [*Λ*]_*k,k*_ = *λ_k_* can be interpreted as spatial frequency of the corresponding structural harmonic (eigenvector) *u_k_*. For each subject, functional data at each timepoint *S_t_* is then projected onto the structural harmonics by assessing spectral coefficients 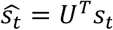, and filtered into two components with ideal low- and high-pass filters (Fig. 1C). A fixed value of *c* = 50 spectral components were chosen, to be common to all acquisitions, and avoid task- or individual-biases that could affect the following classification. The filtering operation yielded a low-frequency functional activity component 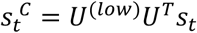, which is coupled to the structure, and a high-frequency one 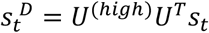, more decoupled from the structure (where *U^(lOW)^* and *U^high^* are *N_ROI_ × N_ROI_* matrices with the *c* first eigenvectors complemented by zeros, and with *c* first columns of zeros followed by the *N_ROI_* – *c* last eigenvectors, respectively). Pairwise Pearson’s correlations of *S^C^* and *S^D^* time courses were computed to obtain c-FC and d-FC matrices, respectively. The L2 norm across time of *S^C^* and *S^D^* yielded instead a general measure of coupling and decoupling for each node, and the ratio between the two corresponds to the SDI (Fig. 1D).

### 2.4 Decoding and fingerprinting networks of the SDI

A two-factor ANOVA with regional SDI values as dependent factor, and subject and task as independent factors was performed to identify brain patterns of task and subject main effects (decoding and fingerprinting patterns, respectively; significant F-values with *p* <.05, accounting for Bonferroni correction across regions). To assess the relative value of SDI and FC nodal strength for task decoding and individual fingerprinting, an additional three-factor (subject, task and measure type; i.e., SDI or FC strength) ANOVA on concatenated regional SDI and FC strength values was performed. The interaction terms [task*measure] and [subject*measure] indicate whether the effect of task or subject on brain patterns depends on the way such patterns are quantified; i.e., structure-function coupling or functional connectivity alone.

### 2.5 Task decoding

Prior to task classification, task paradigms were regressed out from functional time courses to minimize confounds from paradigm-imposed timings, aiming at keeping only differences due to the specific task-related states. Five SVM analyses with *N_BS_* = 8 classes were performed to classify a brain state *bs* (*bs* = 1,…,*N_BS_*; i.e., resting state or one of the 7 tasks) based on the *N_ROI_* × *N_E_ · N_BS_ · N_S_ = 379* × 1600 nodal feature matrices of (1) FC nodal strength and (2) SDI patterns, as well as based on the (*N_ROI_* · (*N_ROI_* – 1) / 2) ×*N_E_* · *N_BS_* · *N_S_* = 71631 × 1600 edgewise feature matrices of (3) FC, (4) c-FC and (5) d-FC values, from all subjects and acquisitions. A 100-fold (*leave-one-subject-out)* cross-validation was implemented, where the *N_E_* · *N_BS_* = 16 acquisitions from one subject were excluded for each training fold and used as test data. For each training-test loop, a *one-versus-one* multiclass linear SVM classifier with error-correcting output codes modelling was trained on standardized training data (i.e., each predictor variable was centred and scaled to unit variance) using the *fitcecoc* MATLAB v.R2019b function and used to predict the task in the test data (Allwein et al., 2000; Furnkranz, 2002).

### 2.6 Individual fingerprinting

A second set of SVM classifications, with the same five sets of features (see paragraph 2.6), but with *N_S_* = 100 classes, was performed to identify individuals based on their functional or structure-function coupling characteristics. Two different classification and cross-validation settings were explored, considering data obtained from matching or discordant tasks: (1) identification of a subject *s* doing a specific task *bs*, based on all other tasks and individuals. This was implemented with a 800-fold (*leave-one-subject’s-task-out*) cross-validation, where the *N_E_* entries (two different encoding directions) of subject *s* doing task *bs* were excluded for each fold; (2) identification of a subject s doing a specific task *bs*, from entries related to only one other different task (all pairwise combinations explored). This was implemented with a set of *leave-one-subject-and-task-out* cross-validation analyses on data subsets including only entries from a specific task and subject in the test fold, and only entries from a specific different task (all subjects, *N_S_* data points) in the training fold. For each training-test loop, a *one-versus-all* multiclass linear SVM classifier with output codes modelling was trained on standardized training data and used to predict the subject in the test data (Allwein et al., 2000).

### 2.7 Multivariate correlation with cognition

PLSC analyses (Krishnan et al., 2011) were performed to assess the presence of multivariate correlation patterns between the five sets of nodal and edgewise brain features and 10 cognitive scores across subjects. For the cognitive scores, the 10 cognitive subdomains tested in the HCP were considered, namely, episodic memory, executive functions, fluid intelligence, language, processing speed, self-regulation/impulsivity, spatial orientation, sustained visual attention, verbal episodic memory and working memory (Barch et al., 2013). For subdomains for which more than one unadjusted raw score was available, a single score was obtained by data projection onto the first component from a principal component analysis (Supplementary Fig. 3). For each brain feature, PLSC was repeated *N_BS_* times, each time considering only brain values (FC nodal strength, SDI, FC, c-FC, or d-FC) obtained during one task. Given the dimensionality of the data, each PLSC analysis outputs 10 pairs of so-called brain-cognitive saliences corresponding to the left and right singular vectors of the data covariance matrix; 10 singular values indicating the amount of explained covariance; and 10 sets of brain and cognitive latent scores corresponding to data projections onto the brain and cognitive saliences, respectively. Statistical significance of multivariate correlation patterns was assessed with permutation testing (1000 permutations) for testing 10 singular values (McIntosh and Lobaugh, 2004; Zöller et al., 2017). Reliability of nonzero salience values was assessed with bootstrapping procedure (1000 random samples) and computing standard scores with respect to the bootstrap distributions (salience values were considered reliable for absolute standard score > 3) (McIntosh and Lobaugh, 2004; Zöller et al., 2017). Moreover, the generalizability of the multivariate correlation patterns obtained with each PLSC analysis was assessed with a 10-fold cross-validation procedure in the following way: first, 10 test subjects were removed from the dataset; second, brain and cognitive saliences were estimated from the remaining data (i.e., from the training set which includes *N_S_* – 10 = 90 subjects); third, test-subject data were projected onto the saliences obtained from the training set to obtain the test latent scores; fourth, the correlation between original and test latent scores was evaluated. In case of generalizable multivariate correlation patterns, one would expect that original and test latent scores align along the identity line (Loukas et al., 2021). Finally, the *r*-squared (squared Pearson’s correlation) between the latent scores was used to quantify the amounts of cognitive traits’ variance explained by the five different brain features. For the edgewise brain features (FC, c-FC, d-FC), a cortical summary of the edgewise saliences was obtained by summing the salience weights of all the edges attached to the individual brain regions.

## 3. Results

### 3.1 Group-level structure-function coupling patterns are consistent across tasks

The structure-function coupling assessed with SDI at the nodal level, and with c-FC and d-FC and the edge level, yielded brain patterns of regional and edgewise values for each subject and run which were consistent across tasks (resting state and seven tasks: emotion, gambling, language, motor, relational, social, working memory; each acquired with 2 phase encoding directions). Average SDI, c-FC, and d-FC profiles across subjects for each state are reported in Supplementary Fig. 1 and Supplementary Fig. 2. Consistently with previous work (Preti and Van De Ville, 2019), we observed relatively strong structure-function nodal coupling (lower SDI) in sensory and particularly in visual areas, and relatively strong nodal decoupling (higher SDI) in high-level cognitive networks (Supplementary Fig. 1). Functional connectivity information extracted from the low spatial frequencies of the structural connectome (c-FC) was qualitatively similar to classical functional connectivity (FC), with strong connectivity within visual and somatosensory networks, and low functional connectivity between the default mode (DMN) and limbic networks, and the other brain circuits. Conversely, functional connectivity matrices obtained from high spatial frequencies of the structural connectome (d-FC) were sparser and displayed both anti-correlation and positive-correlation patterns within and between resting state networks. Subcortical regions mainly showed d-FC anti-correlation patterns with cortical circuits (Supplementary Fig. 2).

### 3.2 SDI task decoding and fingerprinting patterns are spatially distinct

As a first step, we investigated the existence of possibly distinct brain patterns of structurefunction coupling associated with inter-task and inter-individual variability, respectively. To this end, a two-factor ANOVA assessing differences of nodal SDIs across subjects and tasks yielded two spatially distinct whole-brain patterns, characterized by a significant effect for either task or subject. In Fig. 2, nodes with significant *F*-values are visualized as non-zeros, with *p*<.05, Bonferroni-corrected for the number of brain regions. The task decoding pattern (Fig. 2A) clearly involves more prominently regions belonging to unimodal brain circuits, in particular parts of the visual, somatomotor, and auditory networks. On the contrary, the fingerprinting pattern (Fig. 2B) was spatially more distributed, and concerned posterior parietal regions, including fronto-parietal and transmodal cortices that have been consistently reported to contribute to subject identification from functional connectivity (Finn et al., 2015), and to a lesser extent visual, somatomotor, and auditory networks with lower contributions form the anterior DMN and limbic system. Moreover, combined ANOVA analyses including both SDI and FC nodal strength, with subject, task, measure (SDI or FC nodal strength), and first order interactions as explanatory factors, show a significant combined effect of both task and measure (task-measure interaction), and subject and measure (subject-measure interaction) for all brain regions (*p*<.05, Bonferroni-corrected for the number of brain regions), indicating that SDI and FC nodal strength contribute differently to both task and subject identification.

**Fig. 2.**
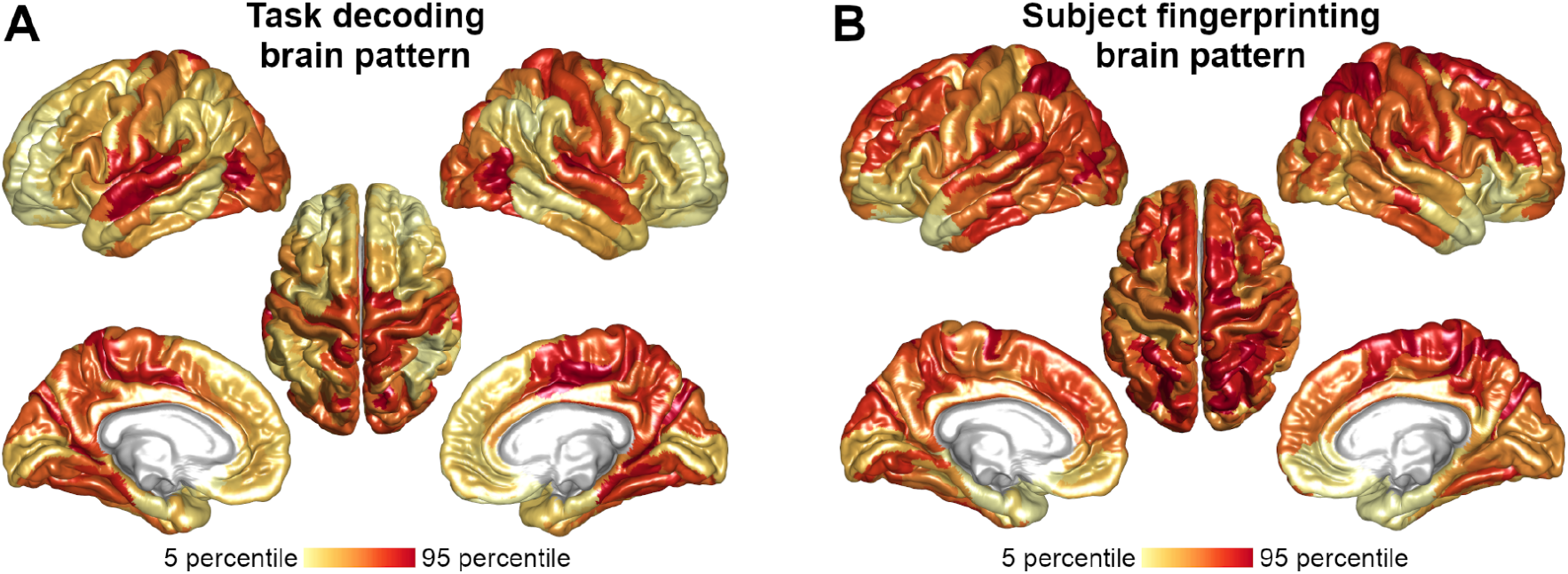
Brain patterns of task (*decoding effect*) and subject (*fingerprinting effect*) main effects on nodal structure-function coupling. Two-Factors ANOVA, significant *F*-values (p<.05 Bonferroni corrected) with colormap scaled between the 5th and 95th percentiles across brain regions (task effect: *F* 20.7-130.0; subject effect: *F* 6.6-15.8, respectively).

### 3.3 Structure-function coupling is able to decode task-related brain states

SVM was used to classify different task-related states (resting state and seven tasks) based on nodewise or edgewise values of functional connectivity as well as structure-function coupling, where task paradigms were regressed out from functional time courses.

For the nodewise metrics, task-classification based on nodal structure-function coupling (SDI) reached an accuracy of 0.703 (chance level = 0.125), higher than the one based on FC nodal strength (0.544), showing that SDI is able to outperform a nodal measure (i.e., with equal dimensionality) based on functional data only (Table 1, first column). When keeping the full dimensionality of connections (71’631 features), accuracy for GSP-derived FC values reached 0.893 for c-FC and 0.873 for d-FC, comparable to conventional FC (0.919), showing that structure-function dependencies alone, analogously to FC, are able to well characterize resting state and the different task conditions.

**Table 1.** Task decoding, subject fingerprinting, and brain-cognition relationships. First column: task decoding accuracies for nodewise (FC nodal strength; SDI) and edgewise (FC, c-FC, d-FC) functional and structure-function coupling measures estimated with 100-fold *leave-one-subject-out* cross-validation and *once-versus-one* multiclass linear SVM classifier. Second column: subject fingerprinting accuracies estimated with 800-fold *leave-one-subject’s-task-out* cross-validation and *one-versus-all* multiclass SVM classifier. Third column: brain-cognition *r*-squared (*r^2^*) computed as the squared Pearson’s correlation coefficient between the brain and cognition latent scores obtained from significant partial least squares correlation (PLSC) components. The brain-cognition *r^2^* quantifies the amount of inter-individual cognitive traits’ variance explained by the five different brain features, respectively.

| | Task Decoding<br>*accuracy* | Subject Fingerprinting<br>*accuracy* | Brain-Cognition<br>$r^2$ |
| --- | --- | --- | --- |
| FC nodal strength | 0.544 | 0.984 | 0.211 |
| nodal SDI | 0.703 | 0.999 | 0.180 |
| FC | 0.919 | 0.964 | 0.224 |
| c-FC | 0.893 | 0.972 | 0.209 |
| d-FC | 0.873 | 1.000 | 0.654 |

### 3.4 Structure-function *de*coupling represents an individual fingerprint of brain organization

In addition to characterizing different task-related states, structure-function coupling measures also revealed to be highly specific to different individuals, which was also the case for functional connectivity. Accuracies for the identification of subjects ranged in fact from 0.964 for edgewise FC to about 1 for nodewise SDI and edgewise d-FC (chance level = 0.010) as assessed with 800-fold *leave-one-subject’s-task-out* cross-validation setting (Table 1). Both nodewise and edgewise structure-function coupling measures performed slightly better than their counterparts based on functional connectivity alone (Table 1). Next, we attempted to identify individuals based on training the SVM classifier on only one task and testing it on another task (all task combinations explored; *leave-one-subject-and-task-out* cross validation). Our results show that even in this more challenging classification setting, subject identification was possible for all functional and structure-function coupling measures, with accuracies largely above chance level (Fig. 3). When considering nodewise measures, fingerprinting accuracies were slightly higher for FC nodal strength compared to SDI in most task combinations (average accuracies = 0.752 / 0.663 for FC strength and SDI, respectively). However, in this same cross-validation scenario, the performance of edgewise metrics was particularly interesting to observe. The best (near-perfect) accuracies, in fact, were reached by the decoupled FC, largely outperforming both conventional FC and, in particular, coupled FC (average accuracies = 0.897 / 0.428 / 0.997 for FC, c-FC and d-FC, respectively). In general, predicting the subject from resting state data (training fold) to task data (test fold), and from any task to resting state data, was slightly more difficult than cross-task prediction, although there was not a particular pairwise task combination consistently outperforming the other task combinations (Fig. 3).

**Fig. 3.**
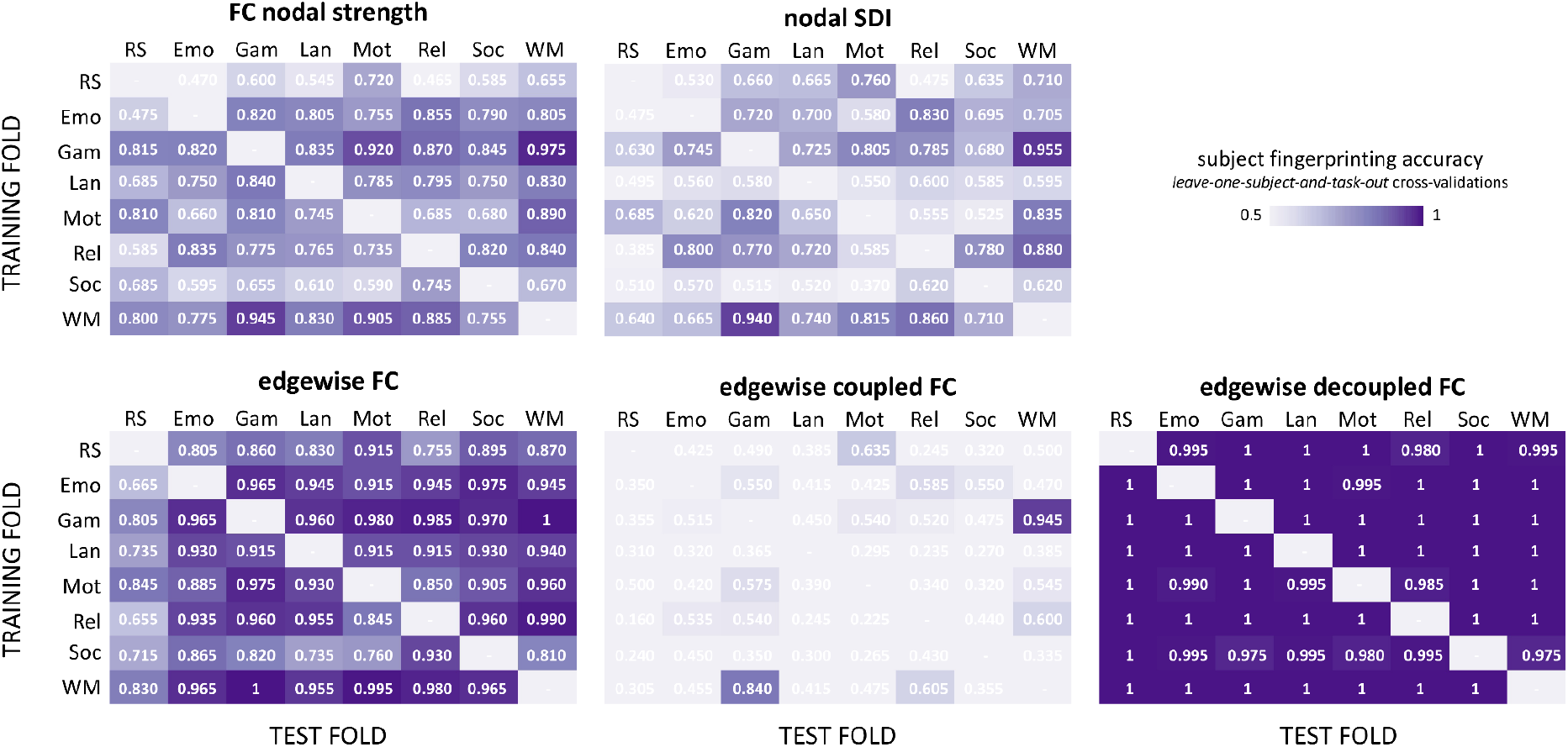
Cross-task fingerprinting accuracies for functional and structure-function coupling measures. Subject classification accuracies when using only one condition -task or resting state-for training (matrices’ rows) and one for testing (matrices’ columns), with all pairwise task combinations explored and for all nodewise (FC nodal strength; SDI) and edgewise (FC, c-FC, d-FC) measures. Classification accuracies were assessed with *leave-one-subject-and-task-out* cross-validation and *one-versus-all* multiclass SVM classifiers. RS=resting state; Emo=emotion; Gam=gambling; Lan=language; Mot=motor; Rel=relational; Soc=social; WM=working memory.

### 3.5 Structure-function *de*coupling explains cognitive traits

Finally, functional and structure-function coupling measures explained inter-individual variations of cognitive traits, particularly sustained attention and fluid intelligence scores. Multivariate correlations between subject-specific brain measures (FC nodal strength, nodal SDI, FC, c-FC, and d-FC) in the different tasks (resting state and seven tasks) and 10 scores measuring cognitive subdomains were assessed with PLSC analyses (one PLSC per task and per brain measure). PLSC identifies linear combinations of brain measures that maximally covary with linear combinations of cognitive scores. PLSC analyses revealed significant multivariate correlation patterns between cognitive traits and all five functional and structurefunction coupling measures mainly during resting state (*p*<.05; Supplementary Table S1). During tasks, brain-cognition multivariate correlations were not statistically significant or statistically significant but weaker compared to resting state, as indicated by lower brain-cognition *r*-squared values (Supplementary Table S1). When comparing the amount of interindividual cognitive traits’ variance explained by the five different brain features, we found that resting state FC nodal strength, nodal SDI, edgewise FC, and c-FC had similar *r*-squared values, ranging from 0.180 for SDI to 0.224 for FC (i.e., 18 to 22%, Table 1). However, edgewise d-FC explained a larger amount of inter-individual cognitive variance, reaching 65% (Table 1). In particular, stronger resting state d-FC in regions belonging to the fronto-parietal network (including the bilateral posterior superior-frontal gyri, dorsolateral frontal cortices, intraparietal sulci, and inferior temporal gyri), and weaker resting state d-FC in somatosensory, limbic and middle temporal regions, were associated with better sustained attention performances, as shown by the d-FC and cognitive saliences that weigh the contribution of individual variables to the overall multivariate pattern (Fig. 4C). Conversely, larger resting state FC nodal strength, SDI, FC, and c-FC specifically related to a cognitive profile characterized by higher fluid intelligence and spatial orientation, and lower, sustained attention and verbal episodic memory scores (Fig. 4A-B and Supplementary Fig. S4). The FC nodal strength, SDI, FC, and c-FC cortical patterns relating to cognition were spatially similar and mainly involved somatosensory, association, and temporo-parietal brain regions. 10-fold cross-validation analyses indicated good brain and cognitive patterns generalizability, with Pearson’s correlation values between original and test latent scores ranging from 0.78 to 0.99 (Supplementary Fig. 5).

**Fig. 4.**
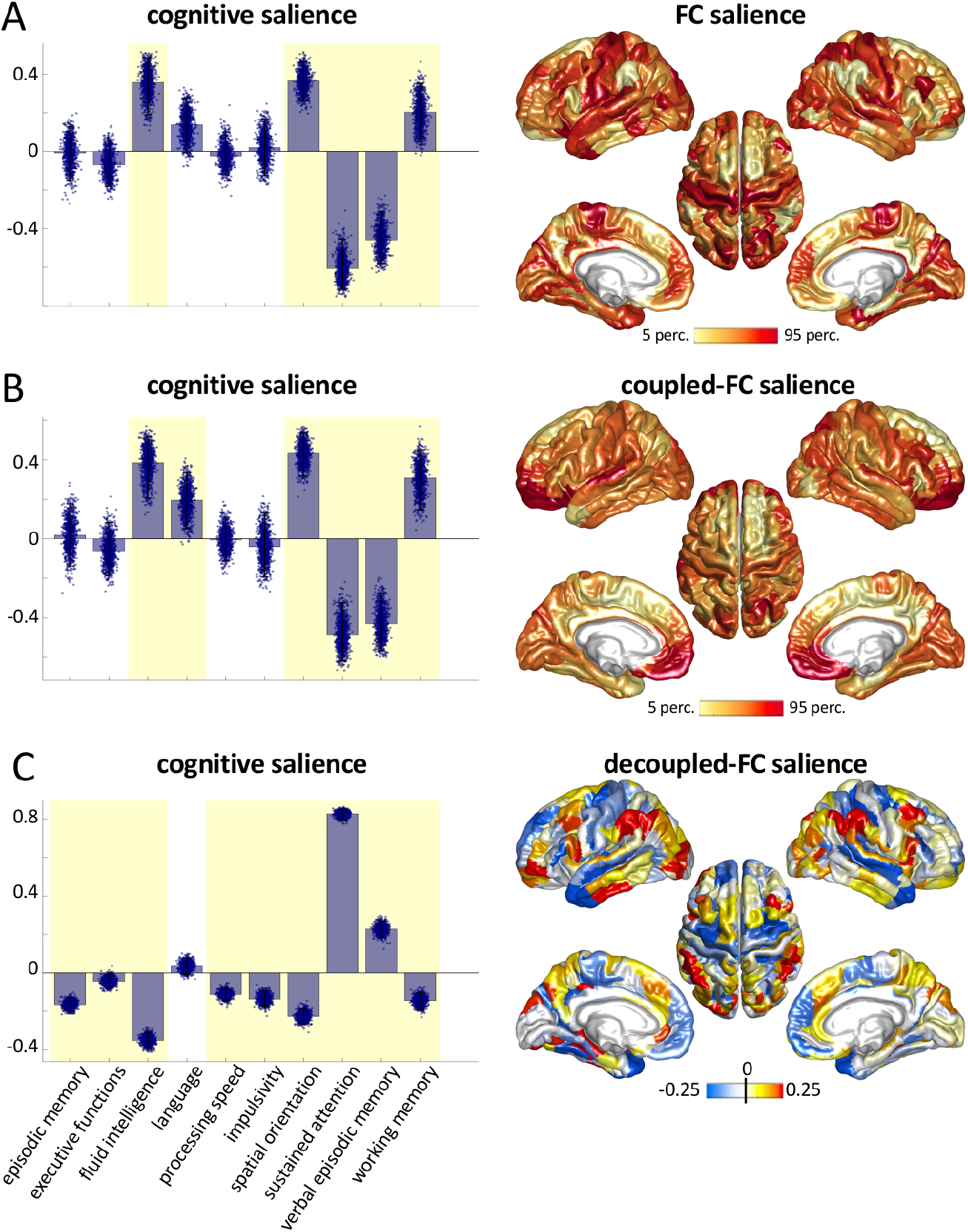
Multivariate correlation patterns between classical, coupled-, and decoupled functional connectivity during rest and cognitive traits. Significant Partial Least Square Correlation (PLSC) patterns between cognitive traits (first column) and resting state FC (A), c-FC (B), and d-FC (C) (second column). First column: cognitive saliences. Bars and single dots represent the salience average and dispersion over 1000 bootstraps; yellow shading indicates cognitive salience weights significantly different from zero. Second column: brain saliences plotted on the cortical surface. For FC and c-FC (A, B) the significant salience weights were positive, as represented by the yellow-to-red colormap. For d-FC (C) the significant salience weights ranged from negative to positive values, as represented by the blue-to-red colormap.

## 4. Discussion

Functional neuroimaging data have shown to provide measures of activity and connectivity with the ability to predict brain states in relation to task execution, as well as to identify individual subjects in a group (Finn et al., 2015; Haynes and Rees, 2006; Richiardi et al., 2011, Van De Ville et al., 2021). In parallel, brain morphology (Wachinger et al., 2015) and structural connectivity (Kumar et al., 2017; Yeh et al., 2016) revealed as well the capability of uniquely identifying individuals. However, brain function and structure are conventionally considered separately and the potential of structure-function coupling in state prediction (task decoding) and subject identification (individual fingerprinting) remains unexplored.

In relation to the first, the way brain function couples to the underlying structure is likely to adapt to the demands of the task. In line with this, task-related functional activity was shown to be well predicted from structure only in selected brain regions, different for each task (Wu et al., 2020). However, how this structure-function relationship depends on external stimulation, cognitive engagement, and affective state, and whether this can be useful to decode different brain states is still an open question (Suárez et al., 2020). Concerning individual fingerprinting, given the high reliability of both structural and functional brain features in subject identification, we could expect structure-function coupling profiles to also uniquely characterize individuals, providing a new dimension of inter-individual differences in brain organization. In line with this hypothesis, a recent study showed that the extent of alignment between structure and function correlates with individual differences in cognitive flexibility (Medaglia et al., 2018).

With these premises, we expanded here previous research by introducing new measures of structure-function coupling at the level of single brain connections (c-FC, d-FC) and by identifying the task decoding and individual fingerprinting potential of such structure-function nodal and edgewise patterns. Specifically, our work shows that the structure-function coupling can predict brain states with high accuracy. The Structural-Decoupling Index and the function connectivity component decoupled from structure (d-FC) revealed able to identify individual subjects in a group with near-perfect accuracy (Table 1), indicating that the pattern of structure-function coupling is an intrinsic feature (or *fingerprint*) of an individual’s brain organization. The idea of a ‘deep’ functional fingerprint, independent from brain state configuration, is consistent with recent works reporting good cross-task subject identification from FC data (Abbas et al., 2020; Amico and Goñi, 2018b; Finn et al., 2015) and moderate state-dependency (compared to high subject-dependency) of functional networks (Gratton et al., 2018). Here, we demonstrate that the way brain function aligns (or misaligns) with the underlying structural connectivity provides additional clues on this functional fingerprint.

Therefore, while it is true that structure-function dependencies are sufficiently different across tasks to allow a reliable decoding of brain states, a strong structure-function individual fingerprint exists independently from the task during which brain function is measured. In fact, this fingerprint appears robust to brain state changes, since even a stringent cross-validation setting with pairwise cross-task predictions delivers high fingerprinting accuracies, in particular related to functional connectivity patterns decoupled from structure. This shows, notably, that a great deal of information specific to the individual is present in the high spatial frequencies of the structural decomposition, that is in the portion of functional signals which is more liberal with respect to brain structure. Notably, this finding could be particularly useful in the context of clinical studies (Itani and Thanou, 2021): the information contained in the high frequencies of the structural connectome, which is shown here to distinguish very well among individuals, could in this case represent features characterizing individual patients and reflecting their specific pathological traits. This consideration is reinforced by the PLSC results which indicate that the FC components decoupled from structure explain a significant percentage of interindividual cognitive traits’ variability, at a level that exceeds the performances of other functional and structure-function coupling measures.

As mentioned, structure-function dependencies also deliver high accuracy (0.75 for SDI, 0.89 for c-FC, and 0.87 for d-FC, against a 0.125 chance-level) when decoding task-related states. It is important to remark here that, having regressed out task paradigms, task decoding can still detect differences due to task, but not “artificially” induced ones, dependent on the paradigm timing, which prevents biases due to task particularities. Recent studies have shown that the cortical macro-scale gradient of structure-function coupling found at rest, opposing primary sensory and association cortices (Preti and Van De Ville, 2019; Vázquez-Rodríguez et al., 2019), can be retrieved from task data as well (Baum et al., 2020; Wu et al., 2020), suggesting similar coupling patterns both in intrinsic (rest) and extrinsic brain states. We can indeed observe the same, when comparing average SDI patterns (across subjects) among task conditions. Nonetheless, specific and non-trivial differences across tasks, not clearly visible at the population level (average maps in Supplementary Fig. 1), exist and allow accurate task decoding.

Contributions of brain regions to task and subject identification are in fact not uniformly distributed across the cortex: two clearly distinct maps were highlighted, one for task decoding and one for individual fingerprinting (see Fig. 1). Interestingly, these two maps group brain regions with distinct structure-function coupling properties. The task pattern mainly involved lower-order regions whose functional activity significantly couples with the structural connectome (Supplementary Fig. 1), including somatomotor, visual and auditory cortices (Preti and Van De Ville, 2019). This means that between-task variations of structure-function coupling mainly occur in regions whose functional activity is on average more constrained by the underlying structure. Conversely, the fingerprinting pattern was spatially more spread and extended to frontal and transmodal association cortices whose functional activity tends to decouple from structure (Preti and Van De Ville, 2019). This difference hints at a neurobiological relevance of the way brain activity (tightly or loosely) couples with the anatomical connectivity substrate, both in regard to the mechanisms underlying brain state reconfiguration across tasks, and to how individual uniqueness is expressed in the brain. In addition, a joint analysis of SDI and FC strength indicated that individual levels of region-wise structure-function coupling and of local functional connectivity strength contribute differently to task and subject identification. At the edge level, structure-function coupling, and particularly the functional connectivity component decoupled from structure, outperforms classical wholebrain functional connectivity in a challenging cross-task subject classification setting. These results suggest that the alignment of function with structure reveals additional information with respect to the functional connectivity alone. Future work should be done to consolidate and extend these considerations, for example by including subject-specific structural connectivity information -a non-trivial operation that would lead to the definition of multiple spectral domains for brain signals, but opens the perspective of incorporating inter-subject structural variability in the analysis of functional brain signatures.

Differently from previous work that mainly focused on fingerprint patterns and single cognitive domains such as fluid intelligence, here we explored multivariate correlations between functional and structure-function coupling features, and multiple cognitive traits. We show that inter-individual variations of (nodewise and edgewise) functional connectivity and local structure-function coupling (SDI) during rest consistently explain traits of complex cognition (fluid intelligence, spatial orientation), executive function (sustained attention) and episodic memory (Moore et al., 2015), resembling descriptions of a general intelligence g-factor previously associated with functional connectivity of the default mode network (S. M. Smith et al., 2015). In particular, a relatively stronger nodal structure-function coupling (lower SDI) was associated with better complex cognition, in line with previous work demonstrating a link between less liberal structure-function alignment during task switching and concomitant cognitive flexibility performances (Medaglia et al., 2018). Nonetheless, relatively weaker nodal structure-function coupling was associated with better executive and memory abilities. It might be that certain brain functions, such as complex reasoning, may benefit from more reliable and consolidated brain communication pathways, possibly expressed in a stronger structurefunction alignment (Finn et al., 2017; Medaglia et al., 2018; Suárez et al., 2020). Other functions, such as verbal learning and retrieval or attention maintenance, may conversely benefit from a less constrained structure-function alignment, a configuration that might predispose the individual to the integration of new information. On this line, stronger functional connectivity components decoupled from structure, mainly in fronto-parietal areas, were strongly associated with better sustained attention performances. While speculative, these considerations and research in this direction, particularly investigating the role of the medium and high frequencies of the structural connectome, may offer a new understanding of cognitive control mechanisms (Lerman-Sinkoff et al., 2017). Furthermore, in our analyses the relationship between brain features and cognitive traits was predominant in the resting condition, suggesting that intrinsic rather than extrinsic brain states might better reflect general cognitive abilities. Meanwhile, this observation does not exclude that temporal fluctuations of structure-function coupling levels during tasks or rest might tap into specific cognitive-behavioral subdomains and hence improve the prediction of task performance or cognitive traits, which is another avenue for future research (Van De Ville et al., 2021).

Finally, the spatial patterns of structure-function coupling relating to cognition presented similarities both with the task decoding map in lower-order somatomotor and association cortices, intrinsically characterized by stronger structure-function coupling, as well as with the fingerprinting map in fronto-parietal regions, characterized by more liberal structure-function coupling (Preti and Van De Ville, 2019) (Fig. 2, 4; Supplementary Fig. 1). Recent work showed that structural and functional connectivity present distinct patterns of inter-individual variance as they relate to cognition (Rasero et al., 2021; Zimmermann et al., 2018). Intriguingly, our results extend these findings identifying in the structure-function coupling a possible link between divergent structural and functional connectivity patterns in predicting behavior. In this respect, both the nodewise SDI and the edgewise d-FC capture inter-subject cognitive variability, but along two different axes. Compared to the functional connectivity component decoupled from structure, the coupled component (c-FC) preserves task- and subject-specific information, but to a lesser extent, showing lower fingerprinting accuracies in the cross-task classification setting and weaker brain-cognition relationship. The coupled functional connectivity component may contain large-scale patterns common to individuals in a group, as suggested by its similarity with the classic functional connectivity organization into well-established resting state networks (Supplementary Fig. 2), while the decoupled component may contain a larger proportion of subject-specific information. A further exploration of the full structural connectome spectrum and of its derived functional connectivity components is warranted.

This study has a number of limitations and possible developments. First, the usage of a grey matter parcellation as opposed to a voxel-based analysis impedes a fine-grained characterization of functional territories that can vary across subjects and tasks (Laumann et al., 2015; Salehi et al., 2020; Wang et al., 2015), with possible impact on the quantification of nodewise and edgewise structure-function coupling features. Nevertheless, a parcellation-based approach facilitates inter-subject comparisons, improves the signal-to-noise ratio of the estimated structural and functional measures, and enables a compact representation of brain fingerprints and decoding patterns. Second, group-level structural connectivity information was used for the computation of GSP-derived metrics. While this choice is convenient since it defines a common spectral domain across subjects and tasks, ways to integrate inter-subject structural variability could be explored in the future. Third, this study does not consider timevarying aspects of structure-function dependency (Cabral et al., 2017; Fukushima et al., 2018; Van De Ville et al., 2021): their exploration in the future might provide insight particularly in relation to task decoding and cognitive control mechanisms. Fourth, our analyses are limited to slow temporal scales accessible with fMRI. Previous studies had attempted brain fingerprinting using electrophysiological recordings (Fraschini et al., 2015; Marcel and Millan, 2007; Sareen et al., 2021), but the link between faster brain dynamics and structural topology remains poorly understood (Finger et al., 2016; Glomb et al., 2020). Future research may address how the hierarchy of structure-function dependencies vary at faster temporal scales, possibly carrying distinct fingerprinting and decoding information. Finally, our multivariate correlation analyses explore possible brain patterns relating to cognitive traits, including bootstrap and cross-validation procedure for generalizability assessment. Nonetheless, feature importance in multivariate predictive models of cognition remains difficult to reliably estimate and different machine learning approaches are under investigation (Tian and Zalesky, 2021).

In conclusion, this work demonstrates that structure-function dependencies quantified both at the level of single brain regions and connections form prominent signatures of individual brains’ organization reflecting cognitive and behavioural correlates, while at the same time preserving task-dependent information. In particular, the high spatial frequencies of the structural connectome may contain relevant subject-specific information which deserves further attention in the future.

## Data and code availability statement

All data used in this study are available through the Human Connectome Project, WU-Minn Consortium. The code to implement the analyses performed in this study will be available upon acceptance at *https://github.com/agriffa/GSP_brain_decode_fingerprint.git*.

## CRediT authorship contribution statement

Alessandra Griffa: Conceptualization, Data analysis, Visualization, Writing—original draft, Writing—review & editing. Enrico Amico: Data preprocessing, Writing—review & editing. Raphaël Liégeois: Data preprocessing, Writing—review & editing. Dimitri Van De Ville: Conceptualization, Supervision, Writing—review & editing. Maria Giulia Preti: Conceptualization, Supervision, Data preprocessing, Data analysis, Visualization, Writing—original draft, Writing—review & editing.

## Acknowledgements

Data were provided by the Human Connectome Project, WU-Minn Consortium (Principal Investigators: David Van Essen and Kamil Ugurbil; 1U54MH091657) funded by the 16 NIH Institutes and Centers that support the NIH Blueprint for Neuroscience Research; and by the McDonnell Center for Systems Neuroscience at Washington University.

AG was supported by the Swiss National Science Foundation (grant number 320030_173153). MGP was supported by the CIBM Center for Biomedical Imaging, a Swiss research center of excellence founded and supported by Lausanne University Hospital (CHUV), University of Lausanne (UNIL), Ecole polytechnique fédérale de Lausanne (EPFL), University of Geneva (UNIGE) and Geneva University Hospitals (HUG). EA acknowledges financial support from the SNSF Ambizione project “Fingerprinting the brain: network science to extract features of cognition, behavior and dysfunction” (grant number PZ00P2_185716). RL was supported by the National Centre of Competence in Research - Evolving Language (grant number 51NF40_180888).

